# An inimitable proprotein convertase subtilisin kexin-9 (PCSK9) cleavage site VFAQ on Spike protein along with furin cleavage site makes SARS-CoV-2 unique

**DOI:** 10.1101/2023.05.04.539453

**Authors:** Medha Pandya, Dhruvam shukla, Sejal Shah, Kajari Das, Sushma Dave, Jayashankar Das

## Abstract

A novel coronavirus (2019-nCoV) or Severe acute respiratory syndrome corona virus 2 (SARS-CoV-2) that affects humans has been discovered in Wuhan, China, in 2019. Its genome has been sequenced, and the genetic data was quickly made public. We discovered a novel proprotein convertase subtilisin kexin-9 (PCSK9) cleavage site in the Spike protein of the 2019-nCoV. The recent research also demonstrates that the previously found proprotein convertase 3 (PC3) or furin cleavage site, which was assumed to be unique, is already present in animal corona viruses. In this article, we suggest that the combination of the both proprotein convertase PC3 cleavage site and the PCSK9 site renders SARS-CoV-2 unique in terms of the pathogenicity, potential functional effects, and implications for the development of antiviral drugs.

## Introduction

The covid 19 pandemic triggered by SARS-CoV-2 (severe acute respiratory syndrome coronavirus 2) came up with a global health emergency. SARS-CoV-2, believed to be transformed from bats, is a single stranded RNA virus having a 29kb genome size. It continuously mutates itself making it more virulent day by day. Though initially it was reported as a pneumonia causing virus in Wuhan, China, later on mortality rate has been increased in spite of several antiviral drugs and plasma treatment. Though, many drugs having anti-viral properties targeting host and viral proteins have been developed to fight SARS-CoV-2 infections, most frequent treatment of antibody to neutralize the SARS-CoV-2 spike glycoprotein have shown good results, still it remains the biggest challenge to come out from this pandemic. The incidence rate continues to rise while the fatality ratio climbed dramatically, making it difficult to entirely manage the virus. This pandemic needs serious concern and requires proper attention with detail structure and properties of SARS-CoV-2 as it behaves differently both in native condition as well as post viral infection. It will aid our understanding towards viral action dynamics and mechanism to the host cell [1]. Number of proteins, growth factors receptors, enzymes are produced as inactive form as precursors. These inactive proteins can be transformed into active proteins by the action of some of the furin like proteases proprotein convertases (PCs). These proteases comprise the family of nine members such as furin (PACE), PC1/3, PC2, PC4, PACE4, PC5/6, and PC7 (LPC, PC8) cleaving precursor proteins at multibasic motifs. Two other family members, SKI-1/S1P and PCSK9, cleave regulator proteins involved in cholesterol and fatty acid homeostasis at nonbasic peptide bonds[2]–[4]. Furin is the most studied enzyme till date due to its importance and ubiquity that cleaves the multi basic consensus motifs R-X-R/K/X-RQ. It expresses in all human tissue except colon carcinoma. It has been also implicated in several malignancy and infectious disease especially bacterial and viral pathogen processing such as ebola virus influenza A, H5N1 (bird flu), flaviviruses etc [5]. Hence, there is a strong need of proteome wide identification for furin substrates and detailed information for the cleavage preferences of furin and in silico studies with suitable pipeline [6]–[8]. It requires computational and experimental model which include multistep in silico pipeline for the identification of the protease substrate in the proteome[9]. Recent studies have reported insertion or deletion of some of the unique sequence in SARS-CoV-2 which was not there in earlier reported coronaviruses (CoVs). ACE and CD 26 receptors present on the host cell are responsible for the interaction with viral spike protein. Spike protein mutation in terms of insertion of furin cleavage site is reported in recent SARS-CoV-2 which is more virulent. Apart from all PCs, PCSK9 is known for cleavage of proteins involved in cholesterol and fatty acid homeostasis at non basic peptide bonds. Catalytic domain is conserved in all the PCs[10]. The C-terminal parts of the PCs have unique peptide signature for the guidance to their secretory pathway destination, it also differs in length and structure such as S1P to the cis-Golgi, furin, PC5B, and PC7 targeted for the TGN region and plasma membrane. PACE4, PC5A, and PCSK9 are there at the cell surface[11]. All the PCs trigger viral envelope fusion with cellular membrane by cleavage of viral glycoprotein, hence can deliver viral genome to host cells. Fusion potential of viral membrane protein can be reduced by the mutational changes that can facilitate preparation of attenuated form of vaccine. Secondly, agents that can prevent viral surface protein cleavage, can block fusion capacity as well as viral replication

### Data retrieval and Phylogenetic analysis of S proteins of SARS-CoV-2

Spike protein sequences of coronaviruses isolated from human and animal species were downloaded from the UniProtKb database [12] (table 1). The multiple sequences of human and animal coronaviruses were aligned in ClustalW. The distance-based neighbor-joining (NJ) phylogenetic tree was then constructed based on protein sequences in MEGA software[13].

**Table I.** Protein sequences of Corona viruses spike protein.

| Coronaviruses | Spike glycoprotein UniProtKb |
| --- | --- |
| Rabbit Coronavirus HKU14 | H9AA34 |
| Canine respiratory Coronavirus | C6ZQJ9 |
| Waterbuck Coronavirus US/OH-WD358/1994 | B7U2N3 |
| Sambar deer Coronavirus US/OH-WD388/1994 | B7U2R5 |
| Sable antelope Coronavirus US/OH1/2003 | A4ZU41 |
| Giraffe Coronavirus US/OH3-TC/2006 | A4ZU52 |
| Murine Coronavirus (strain A59) | P11224 |
| Rat Coronavirus Parker | C6GHS2 |
| Severe acute respiratory syndrome Coronavirus 2 | P0DTC2 |
| Severe acute respiratory syndrome Coronavirus | P59594 |
| Middle East respiratory syndrome-related coronavirus | K9N5Q8 |
| Bat coronavirus | Q3I5J5 |
| Rhinolophus affinis coronavirus | A0A023PUW9 |
| Feline Coronavirus strain | P10033 |
| Porcine transmissible gastroenteritis coronavirus | P27655 |
| Porcine respiratory coronavirus | P27655 |

**TABLE II.**
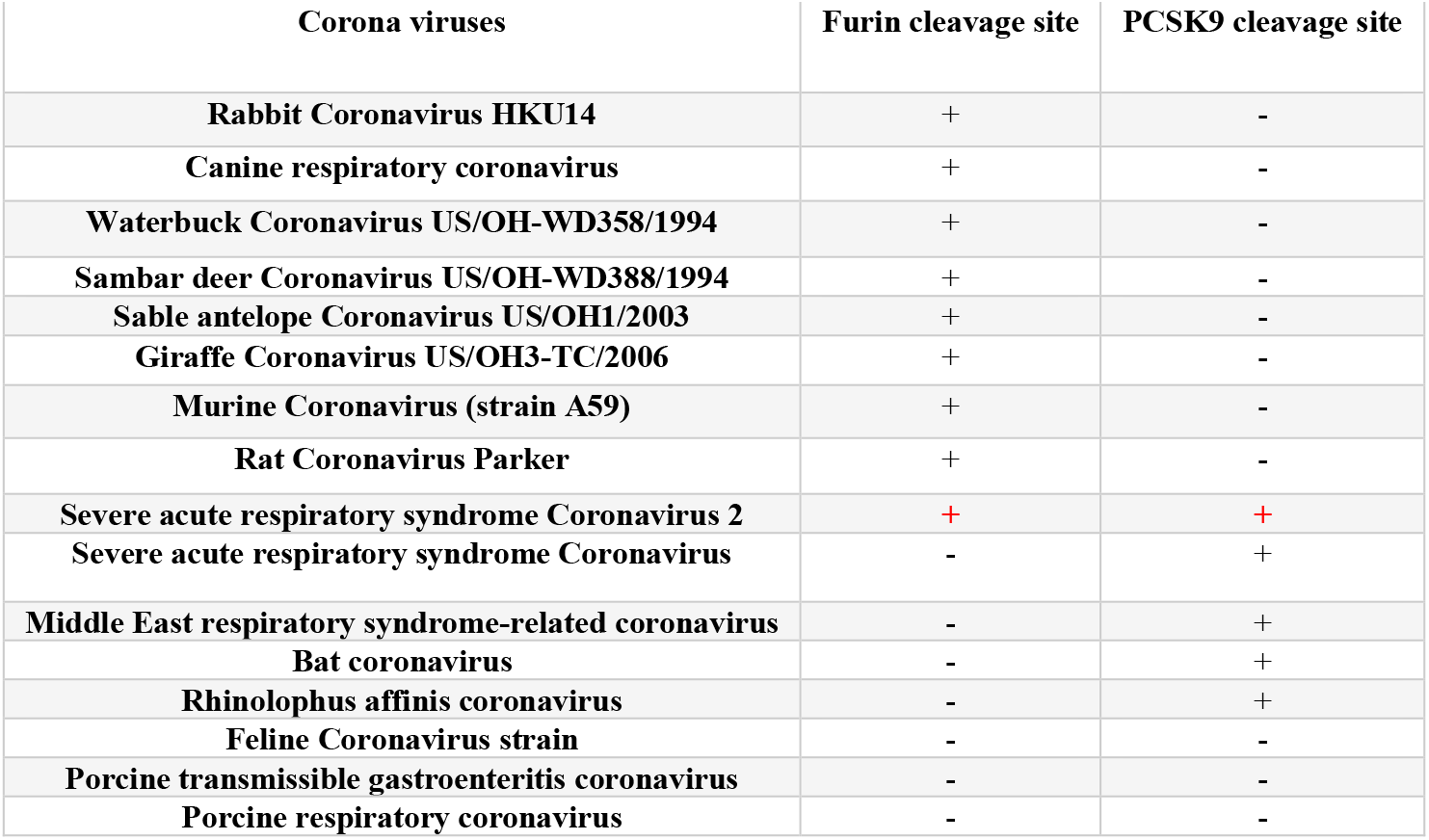
Occurrence of proportion convertase cleavage sites on various coronaviruses spike protein sequence (‘+’ =cleavage site present;’-‘= cleavage site absent)

| Corona viruses | Furin cleavage site | PCSK9 cleavage site |
| --- | --- | --- |
| **Rabbit Coronavirus HKU14** | + | - |
| **Canine respiratory coronavirus** | + | - |
| **Waterbuck Coronavirus US/OH-WD358/1994** | + | - |
| **Sambar deer Coronavirus US/OH-WD388/1994** | + | - |
| **Sable antelope Coronavirus US/OH1/2003** | + | - |
| **Giraffe Coronavirus US/OH3-TC/2006** | + | - |
| **Murine Coronavirus (strain A59)** | + | - |
| **Rat Coronavirus Parker** | + | - |
| **Severe acute respiratory syndrome Coronavirus 2** | + | + |
| **Severe acute respiratory syndrome Coronavirus** | - | + |
| **Middle East respiratory syndrome-related coronavirus** | - | + |
| **Bat coronavirus** | - | + |
| **Rhinolophus affinis coronavirus** | - | + |
| **Feline Coronavirus strain** | - | - |
| **Porcine transmissible gastroenteritis coronavirus** | - | - |
| **Porcine respiratory coronavirus** | - | - |

### Comparison of the SARS-CoV-2 S protein sequence with other corona viruses reveals presence of non-basic PCSK9 cleavage site on SARS-CoV-2

Proprotein convertases; PCs are responsible for activation of many enveloped viruses. PC inhibitors are suggested as promising antiviral drugs for several viruses causing severe infections [10]. The spike protein is a critical target for vaccine development against COVID-19, as it is the key structure on the virus that allows it to enter and infect human cells. By developing vaccines that target the spike protein, scientists hope to provide protection against the virus and ultimately help end the COVID-19 pandemic. Thus a detailed study of PC cleavage sites were the primary focus of the present report. A number of reports show that a variety of viruses exploit a group of host-cell proteases (family of proprotein convertases; PCs), which includes furin, PC2, PC4, PC5, PACE4, PC6 and PC7. Arenaviruses and a few bunyaviruses are reported to be activated by SKI-1/S1P proprotein convertase capable of recognizing a non-basic cleavage site. Like other enveloped viruses that rely on surface glycoproteins for binding and fusion, coronaviruses have the Spike (S) protein, which requires a cleavage by proteases during virion biosynthesis, as well as during entry into target cells. Other proteases, such as the membrane-bound TMPRSS, the lysosomal cathepsins, elastase, and coagulation factor Xa have also been known to be involved in this cycle [14], [15].

To unravel variants of coronaviruses and to understand the possible differences, we accomplish the multiple sequence alignment of the coronaviruses spike protein. The viral spike glycoprotein is indispensable for host cell adhesion via CD26 and ACE2 receptors [16]–[18]. The S protein sequences of ruminants CoVs, Rabbit coronavirus, Canine respiratory coronavirus, Waterbuck coronavirus, Sambar deer coronavirus, and bovine-like Sable antelope coronaviruses, SARS-CoVs, Bat CoVs, MERS CoV were considered in this analysis.PC9 is a newly identified human subtilase that commonly contributes to cholesterol homeostasis [19]. PC9 is mostly located abundantly in plasma and is also expressed in the vasculature of lungs, the primary infection site of SARS-CoV-2. The human PCSK9 gene located on chromosome 1p32.3 is ∼22 kb long and comprises 12 exons encoding a 692 amino acid protein. PCSK9 is expressed mainly in the liver, small intestine, and kidney [20]. No earlier report has yet been published on the potential implication of PCSK9 in SARS-CoV-2 spike processing and viral entry based on in silico study. During this screening process, we discovered presence of four notable amino acid residues **(-VFAQ-)** potentially affecting the cleavage of spike protein by the unique proprotein convertase subtilisin kexin-9 (PCSK9) as our postulate. Along with S protein of SARS-CoV-2, the non-basic cleavage site VFAQ is also speculated to be present on SARS-CoV, Bat CoV, and Rhinopus CoV. In contrast, these unique amino acid residues are absent in Ruminant CoVs and Bovine Covs which is revealed in Fig. 1. In contrast, our result revealed that furin cleavage site is present on Rabbit Coronavirus, Canine respiratory coronavirus, Waterbuck Coronavirus, Sambar deer Coronavirus, Sable antelope Coronavirus, Giraffe Coronavirus, Murine Coronavirus and Parker’s Rat Coronavirus. However, furin cleavage site is absent in SARS-CoV, MERS-CoV and Bat-CoV. The previous study demonstrated by different research groups that insertion of a Furin protease cleavage site in the spike glycoprotein (amino acids 682−689) is strikingly novel in SARS-CoV-2 [21]–[23]. In disparity, our investigation recognises the occurrence of the furin cleavage site assume to be unique for SARS COV-2 already present on the Rabbit Coronavirus HKU14, Canine respiratory coronavirus, Waterbuck Coronavirus US/OH-WD358/1994, Sambar deer Coronavirus US/OH-WD388/1994, Sable antelope Coronavirus US/OH1/2003, Giraffe Coronavirus US/OH3-TC/2006, Murine Coronavirus (strain A59), Rat Coronavirus Parker coronaviruses’ spike proteins as shown in Fig. 2. However, this insertion does not present in other related coronaviruses (SARS-CoV, Bat-CoV, and Rhinopus CoV), but MERS CoV contains a pseudo-binding site. In this article we propose the presence of PC9 cleavage site on novel coronavirus spike protein. Our finding corroborates with the recent wet lab work undertaken by Essalmani et al., (2023) [24].

**Fig. 1.**
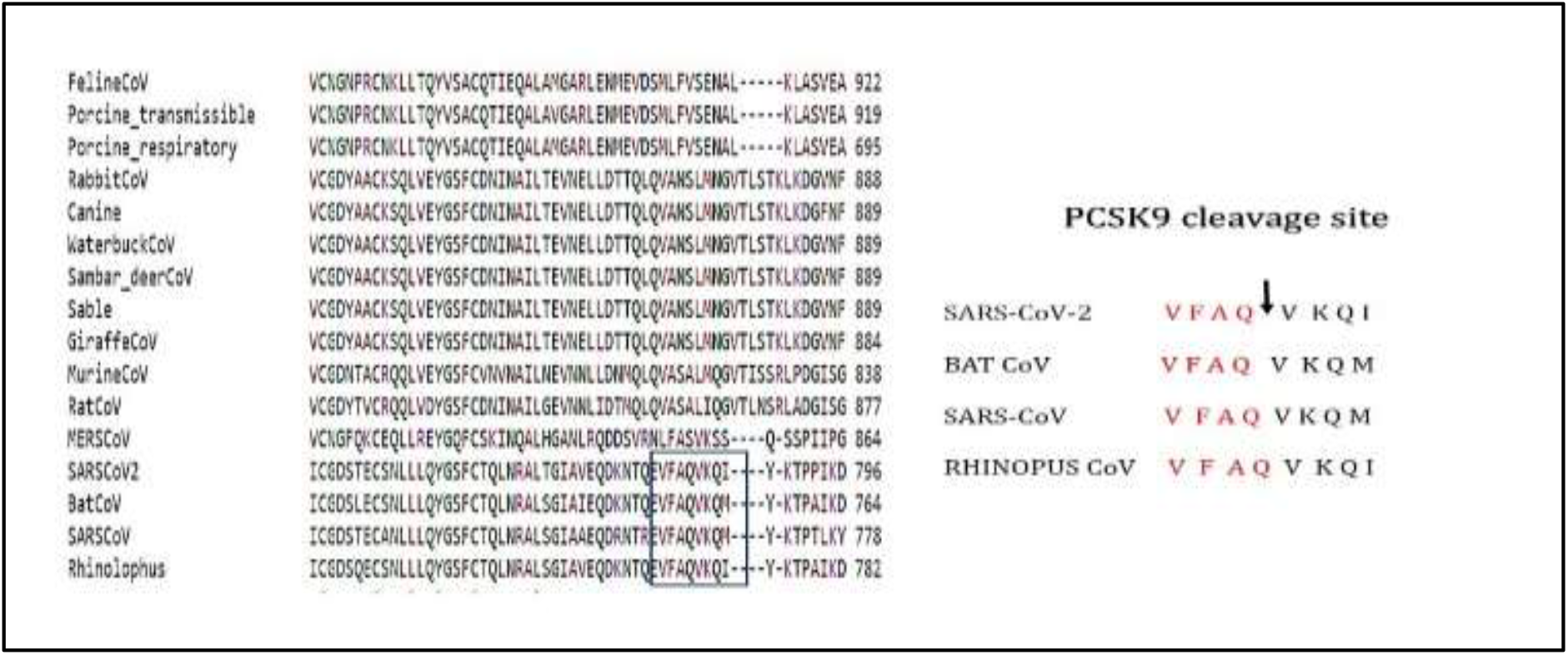
(a.) Multiple sequence alignment of SARS-CoV-2 spike protein with other coronaviruses spike protein. (b) Manifestation of non-basic PCSK9 cleavage sites in SARS-CoV-2, SARS-CoV, Bat Cov and Rhinopus CoVs.

**Fig. 2.**
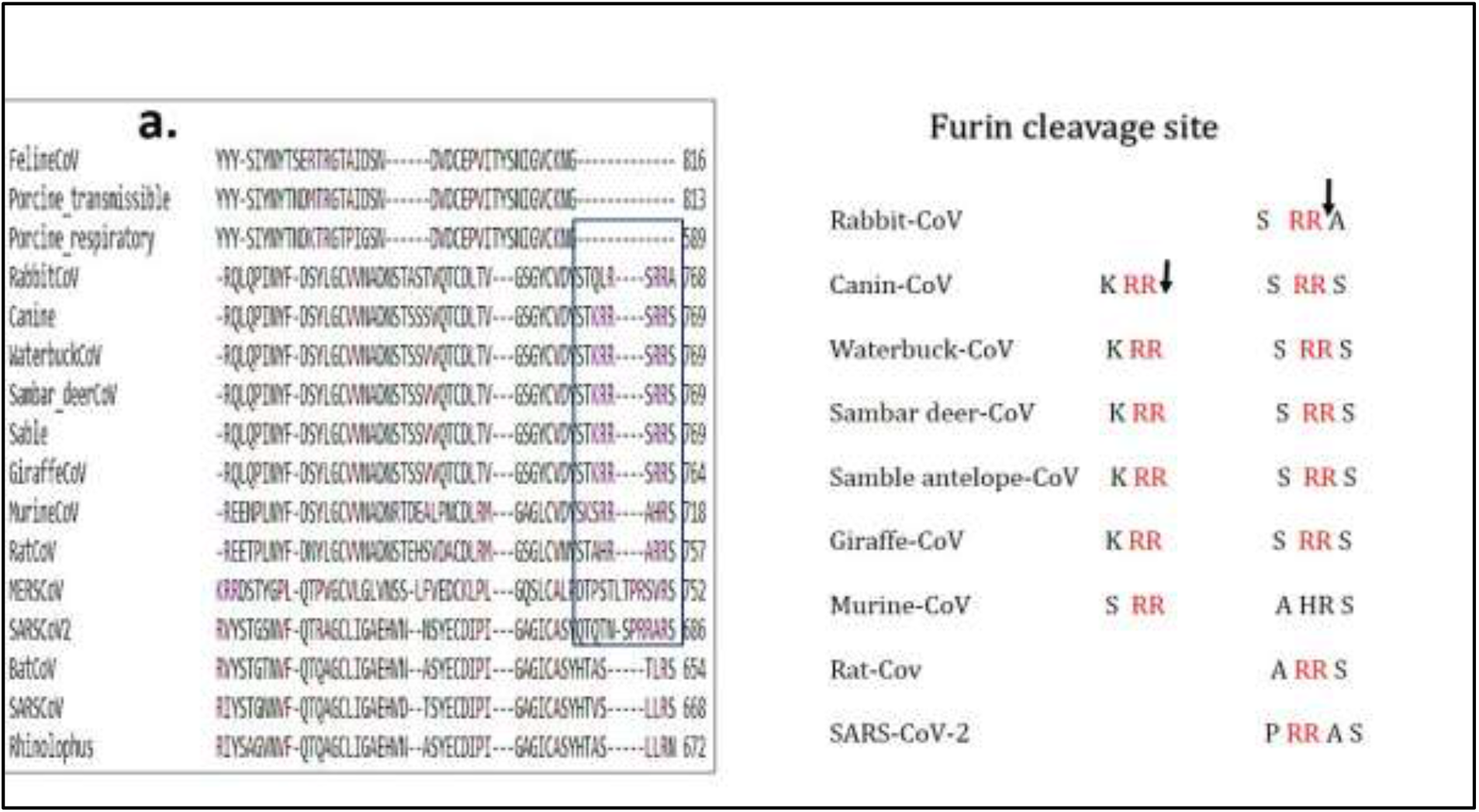
a.) Multiple sequence alignment of SARS-CoV-2 spike protein with other coronaviruses spike protein. (b) Manifestation of polybasic furin cleavage sites in SARS-CoV-2, Rabbit Coronavirus, Canine respiratory CoV, Waterbuck CoV, Sambar deer CoV, antelope CoV, Giraffe CoV, Murine CoV and Rat CoV

PCSK1–7 including furin (PCSK3) cleave their substrates after basic residues, with the typical recognition motif K/R-Xn-K/R↓. In contrast, PCSK8 cleaves after non-basic residues and is best known for its regulation of cholesterol and lipid metabolism. PCSK8 is known to activate transcription factor which is a sterol regulatory element-binding protein [25]. PCSK9 also plays a key role in cholesterol metabolism by directly binding to LDL receptors [26]. The strong binding of PCSK9 with the presence of a non-basic consensus motif could be an indication of this PC’s important role in regulation of viral activation or infection that needs further research. Moreover, the uniqueness of SARS-CoV-2 from other coronaviruses is, it comprises two PC cleavage sites. The polybasic furin cleavage site and non-basic PCSK9 cleavage site in different coronaviruses are visible in table 1. Both the unique PC sites are not found on Porcine CoVs and Feline CoV.

Phylogenetic analysis of corona viruses involves the study of the genetic relationships between different strains of the virus. The phylogenetic analysis helps to understand the evolutionary history of the virus, track its spread, and identify new variants. The genetic material of SARS-CoV-2 is RNA, which mutates at a relatively rapid rate. As a result, the virus can accumulate mutations over time, leading to the development of new strains or variants. The multiple sequence alignment followed by phylogenetic analysis shows the evolutionary history of the virus. In our study, representative S protein sequences of CoVs from human and animal hosts were selected to investigate and compare phylogenetic relationships (Fig.3.). In order to study molecular evolution of spike protein in coronaviruses, the phylogenetic analysis was performed. The phylogenetic tree demonstrates that SARS-CoV-2 is more closely related to SARS-CoV, Bat CoV and Rhinopus CoVs which contain furin cleavage site. The MERS-CoV, Rat corona virus and Murine corona virus are phylogenetically close to SARS-CoV-2 that encompass PCSK9 site.

**Fig. 3.**
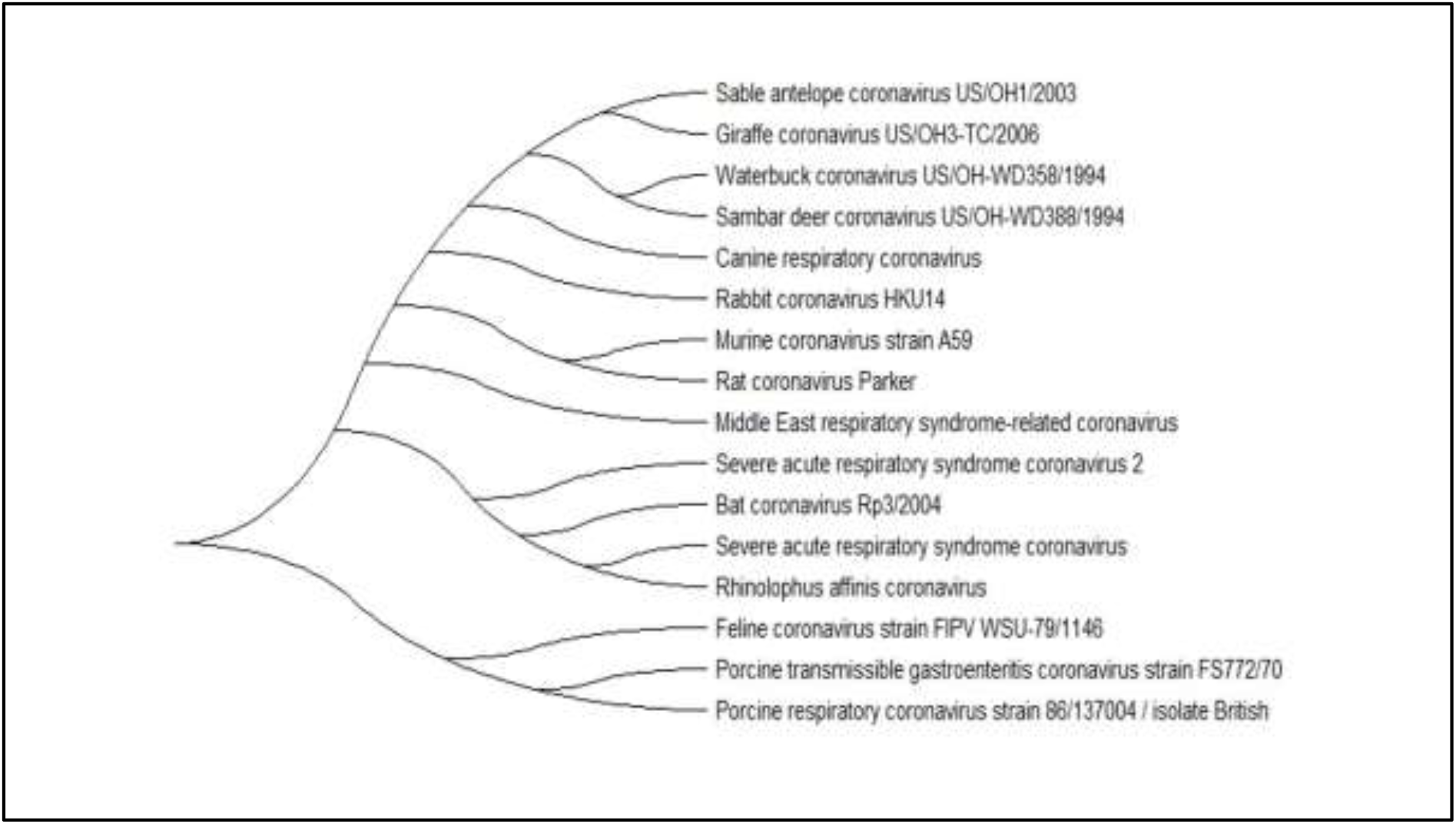
Phylogenetic analysis of amino acid sequence of Spike from various coronaviruses

## Conclusion

In the Spike protein of the 2019-nCoV, we found a unique proprotein convertase subtilisin kexin-9 (PCK-9) cleavage site. The most recent studies also show that animal corona viruses already have the previously discovered proprotein convertase 3 (PC3) or furin cleavage site, which was thought to be unique. In this study, we propose that SARS-CoV-2 is unique in terms of pathogenicity, possible functional effects, and implications for the development of antiviral medicines due to the combination of the proprotein convertase PC3 or furin cleavage site along with the PCSK9 site.

## Acknowledgement

The COVID 19 Omics Research Consortium (CORC) India provided the platform for the research, which is gratefully acknowledged by all authors.

## References

[1] N. Vankadari, “Structure of Furin Protease Binding to SARS-CoV-2 Spike Glycoprotein and Implications for Potential Targets and Virulence,” J Phys Chem Lett, vol. 11, no. 16, pp. 6655–6663, Aug. 2020, doi: 10.1021/acs.jpclett.0c01698.

[2] N. G. Seidah and A. Prat, “The Multifaceted Biology of PCSK9,” Endocr Rev, vol. 43, no. 3, pp. 558–582, May 2022, doi: 10.1210/endrev/bnab035.

[3] N. G. Seidah et al., “The secretory proprotein convertase neural apoptosis-regulated convertase 1 (NARC-1): Liver regeneration and neuronal differentiation,” Proceedings of the National Academy of Sciences, vol. 100, no. 3, pp. 928–933, Feb. 2003, doi: 10.1073/pnas.0335507100.

[4] N. G. Seidah and M. Chrétien, “Proprotein and prohormone convertases: a family of subtilases generating diverse bioactive polypeptides1Published on the World Wide Web on 17 August 1999.1,” Brain Res, vol. 848, no. 1–2, pp. 45–62, Nov. 1999, doi: 10.1016/S0006-8993(99)01909-5.

[5] E. Braun and D. Sauter, “Furin-mediated protein processing in infectious diseases and cancer,” Clin Transl Immunology, vol. 8, no. 8, Jan. 2019, doi: 10.1002/cti2.1073.

[6] M. Dhanalakshmi et al., “Artificial Neural Network-Based Study Predicts GS-441524 as a Potential Inhibitor of SARS-CoV-2 Activator Protein Furin: a Polypharmacology Approach,” Appl Biochem Biotechnol, vol. 194, no. 10, pp. 4511–4529, Oct. 2022, doi: 10.1007/s12010-022-03928-2.

[7] R. Manjunathan et al., “Molecular docking analysis reveals the functional inhibitory effect of Genistein and Quercetin on TMPRSS2: SARS-COV-2 cell entry facilitator spike protein,” BMC Bioinformatics, vol. 23, no. 1, p. 180, Dec. 2022, doi: 10.1186/s12859-022-04724-9.

[8] M. Pandya et al., “Unravelling Vitamin B12 as a potential inhibitor against SARS-CoV-2: A computational approach,” Inform Med Unlocked, vol. 30, p. 100951, 2022, doi: 10.1016/j.imu.2022.100951.

[9] S. A. Shiryaev et al., “High-Resolution Analysis and Functional Mapping of Cleavage Sites and Substrate Proteins of Furin in the Human Proteome,” PLoS One, vol. 8, no. 1, p. e54290, Jan. 2013, doi: 10.1371/journal.pone.0054290.

[10] W. Garten, “Characterization of Proprotein Convertases and Their Involvement in Virus Propagation,” in Activation of Viruses by Host Proteases, Cham: Springer International Publishing, 2018, pp. 205–248. doi: 10.1007/978-3-319-75474-1_9.

[11] S. Duval, “The multifaceted proprotein convertases PC7 and furin: identification of new substrates and physiological relevance,” 2021.

[12] E. Boutet, D. Lieberherr, M. Tognolli, M. Schneider, and A. Bairoch, “UniProtKB/Swiss-Prot,” in Plant Bioinformatics, Totowa, NJ: Humana Press, 2007, pp. 89–112. doi: 10.1007/978-1-59745-535-0_4.

[13] K. Tamura, G. Stecher, and S. Kumar, “MEGA11: Molecular Evolutionary Genetics Analysis Version 11,” Mol Biol Evol, vol. 38, no. 7, pp. 3022–3027, Jun. 2021, doi: 10.1093/molbev/msab120.

[14] N. Seidah et al., “The activation and physiological functions of the proprotein convertases,” Int J Biochem Cell Biol, vol. 40, no. 6–7, pp. 1111–1125, Jun. 2008, doi: 10.1016/j.biocel.2008.01.030.

[15] G. Izaguirre, “The Proteolytic Regulation of Virus Cell Entry by Furin and Other Proprotein Convertases,” Viruses, vol. 11, no. 9, p. 837, Sep. 2019, doi: 10.3390/v11090837.

[16] F. Li, “Receptor Recognition Mechanisms of Coronaviruses: a Decade of Structural Studies,” J Virol, vol. 89, no. 4, pp. 1954–1964, Feb. 2015, doi: 10.1128/JVI.02615-14.

[17] L. K. Gadanec, K. R. McSweeney, T. Qaradakhi, B. Ali, A. Zulli, and V. Apostolopoulos, “Can SARS-CoV-2 Virus Use Multiple Receptors to Enter Host Cells?,” Int J Mol Sci, vol. 22, no. 3, p. 992, Jan. 2021, doi: 10.3390/ijms22030992.

[18] L. Wang and Y. Xiang, “Spike Glycoprotein-Mediated Entry of SARS Coronaviruses,” Viruses, vol. 12, no. 11, p. 1289, Nov. 2020, doi: 10.3390/v12111289.

[19] N. Nassoury et al., “The Cellular Trafficking of the Secretory Proprotein Convertase PCSK9 and Its Dependence on the LDLR,” Traffic, vol. 8, no. 6, pp. 718–732, Jun. 2007, doi: 10.1111/j.1600-0854.2007.00562.x.

[20] G. Lambert, B. Sjouke, B. Choque, J. J. P. Kastelein, and G. K. Hovingh, “The PCSK9 decade,” J Lipid Res, vol. 53, no. 12, pp. 2515–2524, Dec. 2012, doi: 10.1194/jlr.R026658.

[21] N. Vankadari, “Structure of Furin Protease Binding to SARS-CoV-2 Spike Glycoprotein and Implications for Potential Targets and Virulence,” J Phys Chem Lett, vol. 11, no. 16, pp. 6655–6663, Aug. 2020, doi: 10.1021/acs.jpclett.0c01698.

[22] W. Zou et al., “The SARS-CoV-2 Transcriptome and the Dynamics of the S Gene Furin Cleavage Site in Primary Human Airway Epithelia,” mBio, vol. 12, no. 3, Jun. 2021, doi: 10.1128/mBio.01006-21.

[23] A.-B. Butnariu, A. Look, M. Grillo, T. A. Tabish, M. J. McGarvey, and M. Z. I. Pranjol, “SARS-CoV-2–host cell surface interactions and potential antiviral therapies,” Interface Focus, vol. 12, no. 1, Feb. 2022, doi: 10.1098/rsfs.2020.0081.

[24] R. Essalmani et al., “SKI-1/S1P Facilitates SARS-CoV-2 Spike Induced Cell-to-Cell Fusion via Activation of SREBP-2 and Metalloproteases, Whereas PCSK9 Enhances the Degradation of ACE2,” Viruses, vol. 15, no. 2, p. 360, Jan. 2023, doi: 10.3390/v15020360.

[25] N. G. Seidah and A. Prat, “The biology and therapeutic targeting of the proprotein convertases,” Nat Rev Drug Discov, vol. 11, no. 5, pp. 367–383, May 2012, doi: 10.1038/nrd3699.

[26] M. C. McNutt, T. A. Lagace, and J. D. Horton, “Catalytic Activity Is Not Required for Secreted PCSK9 to Reduce Low Density Lipoprotein Receptors in HepG2 Cells,” Journal of Biological Chemistry, vol. 282, no. 29, pp. 20799–20803, Jul. 2007, doi: 10.1074/jbc.C700095200.

